# Increased usage of antiseptics is associated with reduced susceptibility in clinical isolates of *S. aureus*

**DOI:** 10.1101/226738

**Authors:** Katherine Hardy, Katie Sunnucks, Hannah Gil, Sahida Shabir, Eleftheria Trampari, Peter Hawkey, Mark Webber

## Abstract

Hospital acquired infection is a major cause of morbidity and mortality and regimes to prevent infection are crucial in infection control. These include decolonisation of at-risk patients of carriage of MRSA which is commonly achieved by protocols that include the use of chlorhexidine, or octenidine as biocidal agents. There is however no standardised single decolonisation regime agreed upon in the UK or other countries and protocols include a variety of active agents. Antibiotic resistant bacteria cause major problems in hospital medicine and concern has been raised regarding the development of biocide resistance which would cause decolonisation regimes to become unreliable. In this study, we assembled a panel of isolates of *S. aureus* including isolates collected before the development of chlorhexidine and octenidine through to a contemporaneous panel of isolates from a major hospital trust in the UK during a period when the decolonisation regime was altered. We observed significant increases in the MIC and MBC of chlorhexidine in isolates collected from periods of high usage of chlorhexidine. No isolates had a significantly altered MIC or MBC of octenidine apart from those collected after octenidine was introduced into the trust where isolates with four-fold decreases in susceptibility emerged. There was no suggestion of cross-resistance between the two biocidal agents. A combination of VNTR, PCR for *qac* genes and whole genome sequencing was used to type isolates and examine possible mechanisms of resistance. The typing data showed no expansion of a single strain was associated with decreased biocide tolerance and isolates with increased chlorhexidine MIC and MBCs were found from different clonal complexes; CC8, CC22 and CC30. Biocide susceptibility did not correlate with carriage of *qac* efflux pump genes – carriage of *qacA* and *qacB* was detected but, with one exception was restricted to isolates of CC8. Analysis of genome sequence data for closely related pairs of strains with differential biocide susceptibility revealed no common mutations or carriage of accessory elements that correlated with biocide tolerance. Mutations with the NorA or NorB efflux pumps, previously associated with chlorhexidine export were identified suggesting this may be an important mechanism of biocide tolerance. The clinical relevance of decreased biocide tolerance in terms of efficacy of decolonisation therapies remains to be established but we present evidence here that isolates are evolving in the face of biocide challenge in patients and that changes to decolonisation regimes are reflected in changes in susceptibility of isolates. More work is needed to assess the impact of these changes to ensure effective and robust decolonisation protocols remain in place.

## Introduction

Antiseptics, and especially chlorhexidine, have been used widely as one of the key measures implemented in the control of infections caused by methicillin resistant *Staphylococcus aureus* (MRSA) in hospitals. The most widely used approach has been to use antiseptics as part of a decolonisation protocol for patients who are known to be colonised with MRSA in conjunction with nasal mupirocin. However, in some units, especially intensive care units, antiseptics have been used universally for washing of all patients (Climo *et al*., 2013).

Chlorhexidine, which was first synthesised in 1954 (Davies *et al*., 1954), is the most widely used antiseptic. It is a cationic biguanide agent that acts by disrupting the bacterial cell membrane. A more recent introduction to clinical practice is Octenidine dihydrochloride, which was synthesised over 30 years later (Sedlock and Bailey, 1985). Like chlorhexidine it is a cationic biguanide agent that has a broad spectrum of activity.

Due to the widespread usage of chlorhexidine, questions have been raised over the development of resistance (Wesgate et al., 2016., Oggioni et al., 2013). Determination of chlorhexidine susceptibility has most often relied on data from minimum inhibitory concentration (MIC) determinations. The MIC is however an imperfect method for testing biocide resistance where the lethal rather than inhibitory concentration of the agent is of primary importance and needs to be measured. In addition, there is no agreed standardised testing methodology and no national or international agreement regarding an appropriate cut off for defining chlorhexidine resistance. However, many studies have often defined isolates with a chlorhexidine MIC ≥4mg/l as resistant (Horner et al., 2012). No definition has been proposed for Octenidine.

The mechanisms of action and resistance to biocides including chlorhexidine remain poorly understood, although there are several genes that encode efflux pumps which have been shown to be able to influence biocide susceptibility. Of these, *qacA* is the most commonly associated with reduced susceptibility to chlorhexidine in Staphylococci. However, the presence of *qacA* does not necessarily result in expression of resistance to chlorhexidine and conversely resistance can be expressed without the presence of *qacA* (McGann et al., 2011; Longtin et al., 2011).

Most studies that have investigated reduced susceptibility to biocides have studied chlorhexidine. The limited number of studies that have investigated octenidine have looked at the clinical efficacy and have not included susceptibility testing as part of their studies, with the only *in vitro* study failing to select for resistance to octenidine dihydrochloride (Al Doori *et al*., 2007; Spencer *et al*., 2013; Krishna *et al*., 2010). Unlike octenidine, reduced susceptibility has been described in chlorhexidine, with the prevalence rates varying between studies (Horner et al., 2012). Few longitudinal studies have been reported, but one in Taiwan observed an increase in the percentage of MRSA strains with reduced susceptibility to chlorhexidine from 1.7% to 46.7% from 1990 to 2005 (Wang *et al*., 2008). Whilst significant increases in the MIC between both MSSA and MRSA isolates from 1989 compared to 2000 were observed by Lambert (Lambert, 2004). Warren *et al*., observed a non-linear increase in the presence of *qacA/B* positive isolates, markedly increasing in years five and six of their study, but then decreasing in the following two years (Warren et al., 2016). There have also been case reports of the selection of isolates with an increase in chlorhexidine MIC in patients receiving chlorhexidine as part of a decolonisation regime (Johnson et al., 2015).

Despite the reporting of isolates with reduced antiseptic susceptibility, the clinical impact of this is unclear with antiseptics being used at much higher concentrations than the typical MIC or MBC of these isolates. However, there are reports of clinical failures of decolonisation, including both an outbreak of a specific clone of MRSA that had decreased susceptibility to chlorhexidine and the persistence of a hospital clone that carried *qacA* and out-competed a non *qacA* carrying clone (Kampf et al., 2016; Otter et al., 2013; Batra *et al*., 2010).

This study aimed to investigate whether susceptibility to the two most commonly used biocides for the decolonisation of patients colonised with MRSA, chlorhexidine and octenidine dihydrochloride, varied in a unique panel of *S. aureus* strains isolated over an extended period where chlorhexidine use increased and octenidine was introduced.

## Materials and Methods

A panel of 157 *S. aureus* and methicillin resistant *S. aureus* strains isolated between 1928 and 2014 were included in the study (Table 1). This panel includes some of the earliest MRSA isolates from the 1960s and 1970s, from the National Culture Type Collection (NCTC). All MRSA isolates from 2002 onwards were collected from one large NHS trust in the West Midlands of the UK. Since 2002 the hospital has had a policy of prescribing a 5 day course of chlorhexidine to decolonise all known MRSA positive patients. In 2014 universal washing of all patients with octenidine dihydrochloride for the duration of their hospital stay was implemented, with those patients identified as being colonised/infected with MRSA continuing to receive chlorhexidine.

**Table 1:** Number of MRSA and MSSA isolates included in each period and corresponding biocide usage.

| Years |  | (1) 1928-1953 | (2) 1954-2001 | (3) 2002-2012 | (4) 2013-2014 |
| --- | --- | --- | --- | --- | --- |
| **Biocide Usage** | Chlorhexidine | None | Minimal | Significant | Significant |
|  | Octenidine | None | None | None | Significant |
| **Isolates** | MSSA | 15 | 9 | 1 | 0 |
|  | MRSA | 0 | 54 | 52 | 26 |

In this study both screening and clinical isolates were included. The isolates were grouped into four panels (Table 1) to reflect the usage of octenidine dihydrochloride and chlorhexidine from 1928 to 2014 (Groups 1–4). The first group comprised 15 MSSA isolates from 1928–1953 where there was no use of either antiseptic and provides historical context for susceptibility of populations unexposed to antiseptics. The second group contained 54 MRSA and 9 MSSA isolates from 1954–2001 where there was low usage of chlorhexidine and no usage of octenidine. The third group contained 1 MSSA and 52 MRSA isolates from 2002–2012 where chlorhexidine usage was high but octenidine was not used and the final group contained 26 MRSA isolates from 2013–2014 where there was high use of chlorhexidine and octenidine had been introduced. The dominance of MRSA over MSSA isolates reflects routine surveillance practice where MRSA are actively identified in patients on admission but MSSA are not.

The minimum inhibitory concentrations (MIC) of each agent were determined using broth microdilution (following recommendations from EUCAST) ranging in concentration from 0.0029μg/ml to 3μg/ml for octenidine dihydrochloride and 0.0625μg/ml to 64μg/ml for chlorhexidine digluconate. Minimum bactericidal concentrations (MBC) were subsequently determined by inoculation of 10 μl of suspensions following determinations of MICs onto LB agar and observing growth after overnight incubation. The presence of qacA/B was determined in all samples using mutliplex PCR (Vali *et al*., 2008).

A panel of 99/157 isolates which included all *qacA/B* positive isolates and all isolates from groups 3 and 4 were epidemiologically typed using variable number tandem repeats as previously described (Hardy et al., 2006). A selection of 16 isolates from groups 3 and 4 to include the main circulating clones were also genome sequenced using Illumnina paired end sequencing. Reads were then analysed using the ‘nullarbor’ pipeline (v1.2; Seemann et al., 2017) using a standard virtual machine on the MRC CLIMB framework. Pan genomes were generated using ‘roary’ (v8.0), SNPs called with ‘snippy’ (v3.0) and antibiotic resistance genes and mutations identified using ‘ARIBA’ (v2.8.1). Trees were visualised with ‘Phandango’. All packages used default parameters unless stated otherwise.

Data was obtained for the number of packs of chlorhexidine and octenidine dihydrochloride issued by pharmacy in the large teaching Trust from 2008 to 2014 where the MRSA isolates were obtained from. Statistical analysis of changes in susceptibility patterns between groups used chi-squared and Mann-Whitney tests.

## Results

### Usage of chlorhexidine and octenidine

The use of chlorhexidine and octenidine was analysed in our hospital showing a decrease in usage of chlorhexidine and an increase in octenidine use across the study period. The usage of chlorhexidine decreased from 7,061 packs being dispensed in 2009 to 5091 in 2014. In contrast, the usage of octenidine dihydrochloride has increased markedly, with none being prescribed in or before 2013, increasing to 18,844 bottles in 2014.

### Biocide susceptibility of isolates

The MIC and MBC of chlorhexidine was significantly different between the four groups. Chlorhexidine MICs ranged from 0.5 μg/ml to 32 μg/ml, with a significant increase in the mean MICs over time from group 1 to 3, with then a slight reduction in group 4 compared to group 3 (Figure 1). This increase in mean MIC was a result of a shift of susceptibility of most isolates in the population requiring higher MICs rather than being a result of a small sub-population of highly resistant isolates. The MBCs of chlorhexidine showed a similar pattern, the percentage of isolates with an MBC >32 μg/ml increased from group 1 to 3, with 0, 5.5 and 36.5% of isolates having an MBC >32 μg/ml in groups 1, 2 and 3 respectively (Figure 3). The differences in MIC and MBC distributions between the groups were not likely to be random according to statistical tests (chi-squared and Mann-Whitney tests both returned p values <0.05 comparing groups 3 and 4 and 1 and 2).

**Figure 1.**
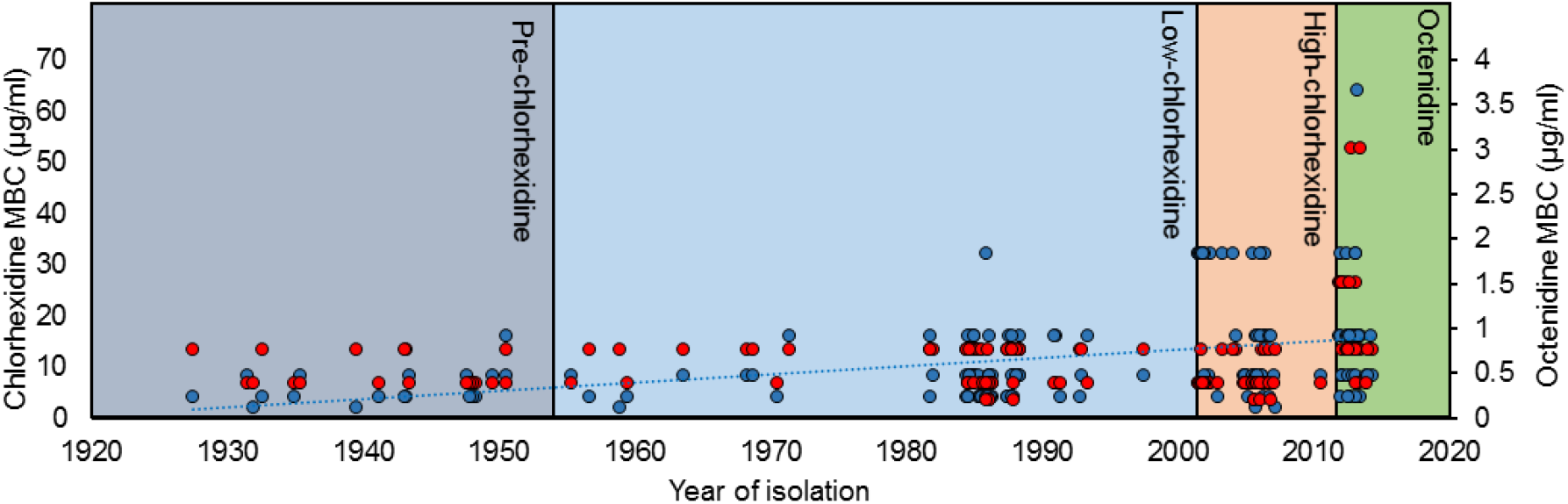
Timeline of mean MBC of chlorhexidine (blue circles) and octenidine (red circles) against isolates. The shaded boxes represent different periods of biocide usage. A trendline (blue, linear) is shown for chlorhexidine but not octenidine where isolates with decreased susceptibility only emerged in the final period.

The MIC and MBC of octenidine dihydrochloride were both lower than chlorhexidine and ranged from 0.09375 μg/ml to 1.5 μg/ml. The MBC of octenidine was stable against isolates in groups 1-3 with all isolates being inhibited by 0.0375-0.75 μg/ml. Isolates with a markedly decreased susceptibility were however isolated in group 4 after introduction of this agent (Figures 1 and 2). The difference between MIC and MBC values for the final group (4) and other groups was statistically significant (P <0.05).

**Figure 2.**
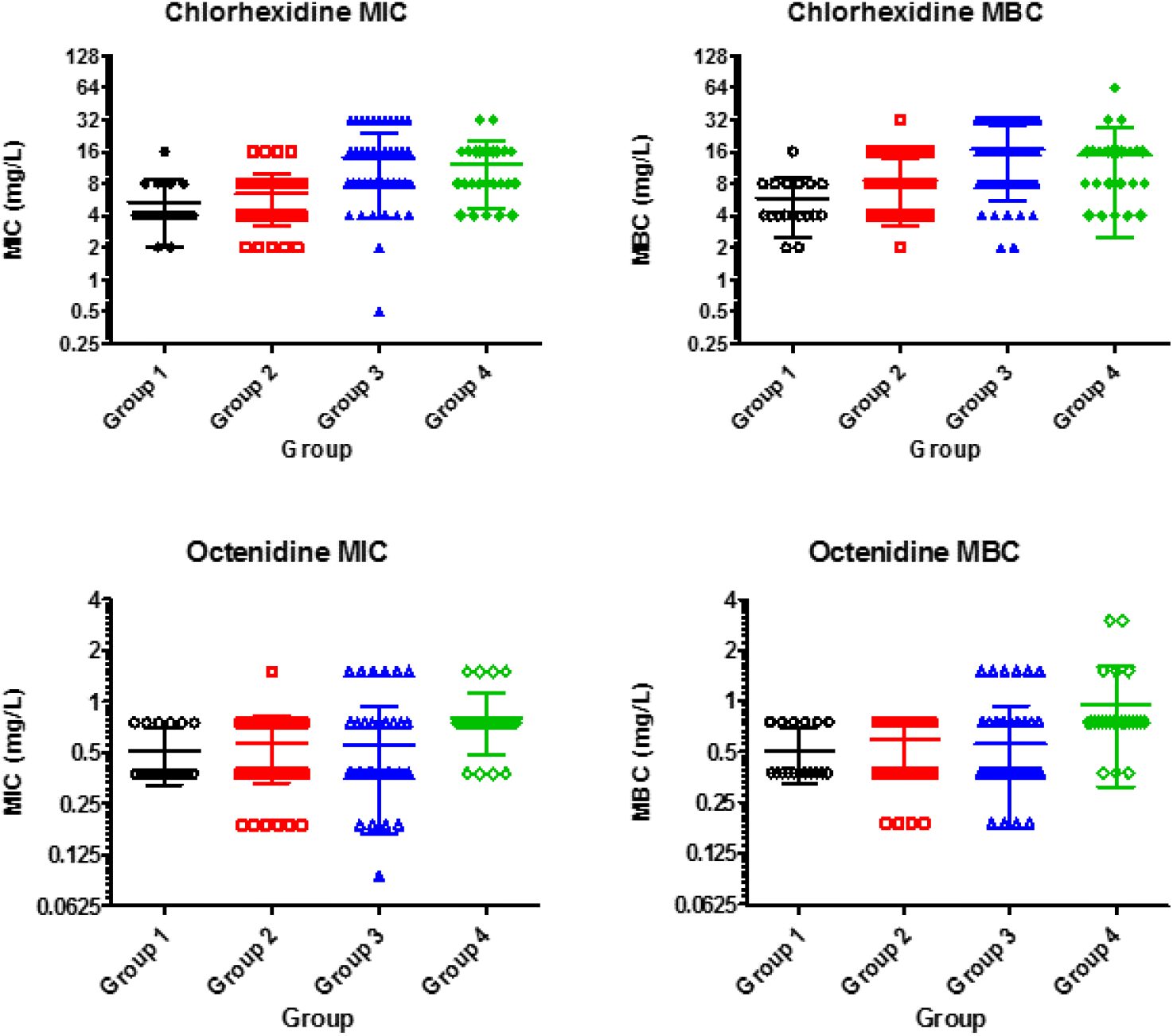
Mean MIC and MBC of *S. aureus* isolates to chlorhexidine and octenidine.

Interestingly, there was no correlation between susceptibility to the two agents, i.e. a raised chlorhexidine MIC was not likely to be reflected by a raised octendine dihydrochloride MIC in the same isolate (tested using Pearson’s correlation test).

The changes in susceptibilty to both agents reflected usage data although it was not possible to statistically analyse these changes.

### Typing of isolates

VNTR analysis of the isolates revealed CC22 to be the most predominant clonal complex, which is the endemic strain within the UK. The other two clonal complexes represented were CC36 and CC8 (Figure 3).

**Figure 3.**
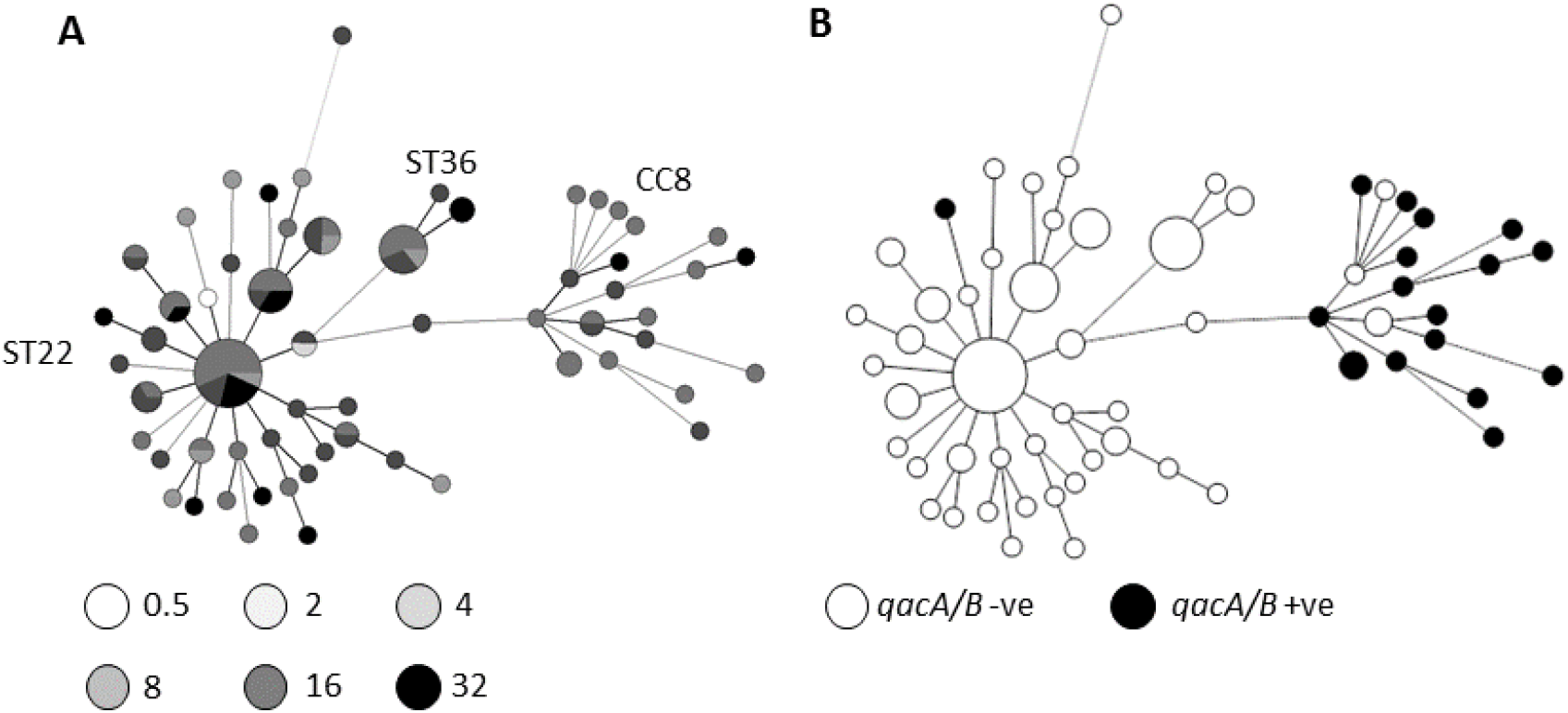
A minimum spanning tree of isolates based in VNTR profile. Sizes of circles reflect number of isolates Panel **A** shows isolates shaded by MIC of chlorhexidine (μg/ml) as per key below the tree. Panel **B** shows isolates found to carry *qacA/B* (black circles).

The phylogenetic analysis failed to identify a specific clone or lineage which showed reduced susceptibility to chlorhexidine (Figure 3). Interestingly, overlaying susceptibility data onto the phylogeny demonstrated several occasions where isolates with the same VNTR profile differed in MIC for both chlorhexidine and octenidine dihydrochloride suggesting the acquisition of decreased susceptibility is not restricted to one clonal complex and that it may be able to evolve independently from various strains (Figure 3).

### Carriage of *qac* genes

A total of 18/157 (11.4%) of all isolates were positive for carriage of the *qacA/B* gene, all of these were MRSA isolates apart from one MSSA isolate. All of the isolates carrying the *qacA/B* gene had a chlorhexidine MBC >4μg/ml. The majority of isolates in he collection with the highest MICs and MBCs did not however carry *qacA/B*. The highest percentage of isolates carrying the *qacA/B* gene were in group 2 (20.6%) as opposed to groups 3 and 4 where only 5.6% and 7.7% of isolates were carrying *qacA/B, qacA/B* was not detected in any of the pre 1954 isolates in group 1. VNTR typing of all the *qacA/B* positive isolates revealed all but one of the *qacA/B* positive isolates clustered and belonged to CC8 (Figure 3). The other isolate was from the ST22 cluster and no isolates from the ST36 cluster carried a *qac* gene.

### Genomic analysis of ST22 isolates

Sixteen strains with related VNTR profiles from groups 3 and four and a range of chlorhexidine MBCs were whole genome sequenced and mechanisms which may contribute to decreased susceptiblity to chlorhexidine were identified. Figure 4 shows the phylogeny of these strains based on a whole genome alignment (produced by ROARY). Comparisons of the accessory genome content, resistance genes (by ARIBA) and presence and absence of core genes attempted to identify common changes in those isolates with highest MBCs. There were no common accessory genes identified or carriage of a known resistance mechanism that correlates with biocide resistance. To further try and identify a tolerance mechanism two pairs of strains with four fold differences in susceptibilty (by MBC) but which were very closely related according to the phlogeny were compared for changes (strains 3 vs 7 and strain 2 vs the reference ST22; HO_5096_041, Figure 5). Analysis of these strain pairs found a small number of SNPs between each (5 vs 20 SNPS between each pair, respectively), the two more resistant isolates both had changes in either of two homologous efflux pump systems; NorA and NorB. Both have been shown to export multiple agents including biocides (DeMarco et al., 2007). Strain 2 carried a SNP within *norB* that results in a change of the NorB protein of Met314Ile. This substitution is adjacent to the predicted translocation pore and may alter the capacity of this strain to export chlorhexidine compared to the reference. Strain 7 had a wild-type *norB* allele but carried a SNP within *norA* resulting in a change in NorA of Ala362Val. Substitution at this site has previously been shown to improve efflux capacity of the protein for some substrates (Kaatz *et al*., 1993). None of the other sequenced strains had changes within either system or in their known regulators.

**Figure 4.**
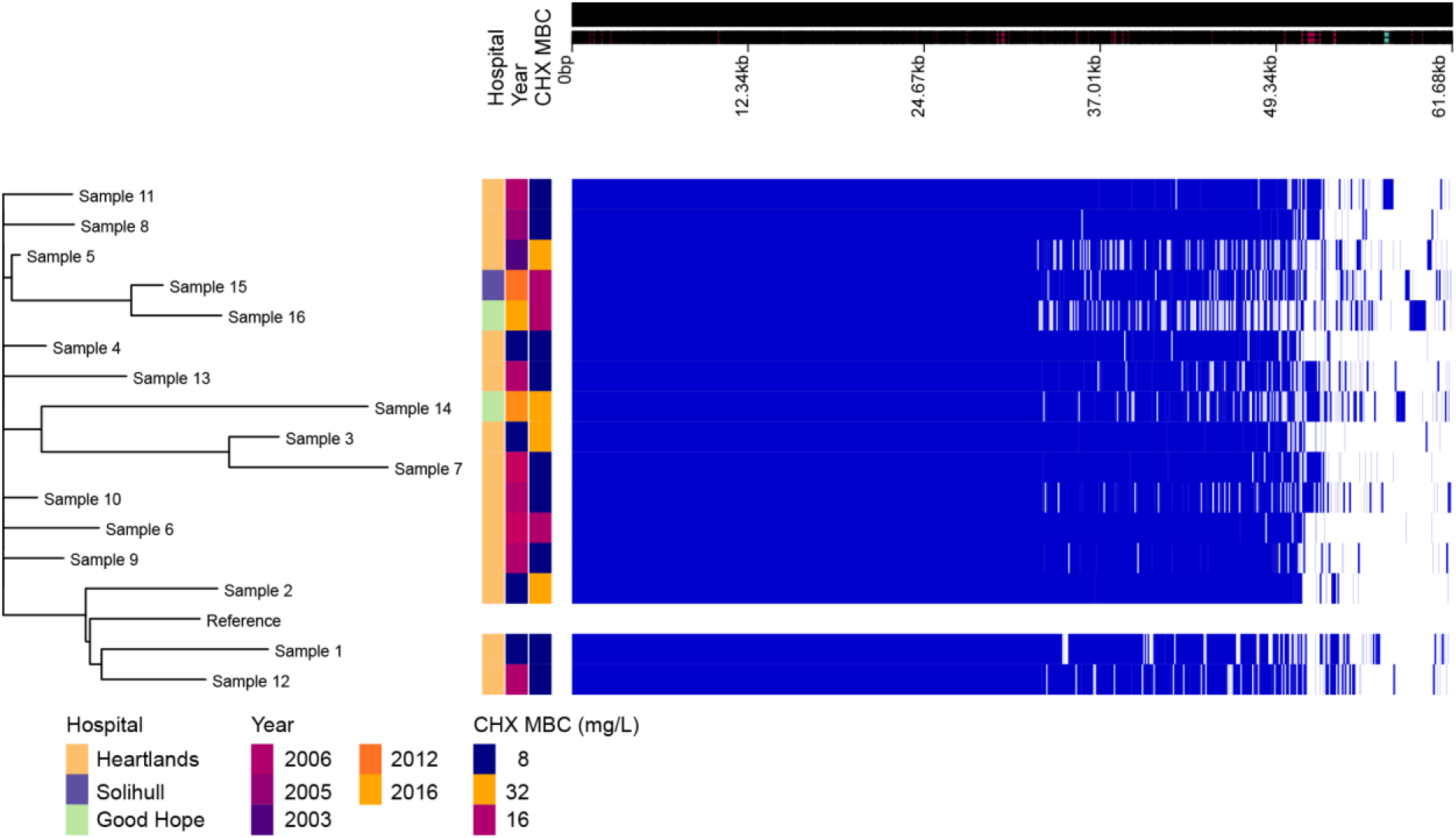
Visualisation of pan-genome analysis by ROARY of 16 isolates. Hospital and year of isolation are indicated as well as chlorhexidine MBC.

## Discussion

This study has highlighted that the increasing use of chlorhexidine and octenidine dihydrochloride is associated with emergence of reduced susceptibility to each agent in *S. aureus*. As described in previous studies (Wang *et al*., 2008; Liu *et al*., 2015) the number of *S. aureus* and especially MRSA isolates demonstrating reduced susceptibility to chlorhexidine increased over time, which in the current study was most marked when the MRSA epidemic was at its peak within the UK.

There have only been a limited number of studies that have investigated reduced susceptibility to chlorhexidine longitudinally and all have been over shorter time periods. Historical isolates included in this study from a period of no antiseptic usage had very low MIC and MBC and increases were then observed in groups two and three. Interestingly unlike other longitudinal studies where there have been similar observations we did observe a reduction in the MICs in the fourth time period. This can be correlated with a reduction in the number of MRSA infections within the UK and reduction in chlorhexidine usage in our hospital.

We also demonstrate a reduction in susceptibility to octenidine following universal introduction. There have been a limited number of studies investigating the clinical efficacy of octenidine dihydrochloride which whilst being small demonstrated comparable efficacy to chlorhexidine, but did not include susceptibility testing as part of the studies (Spencer *et al*., 2013; Krishna *et al*., 2010). The only *in vitro* study failed to select for resistance to octenidine dihydrochloride (Al Doori *et al*., 2007).

Whilst there has been a demonstrable reduction in susceptibility in isolates in the current study and this markedly occurred after the introduction of octenidine into practice the MIC and MBC are still relatively low and significantly below the concentrations that the antiseptics are used at in practice. It is unclear what impact these relatively small changes in tolerance as measured in vitro have in practice. Comparing the MIC values to in use concentrations suggests that it is unlikely that these isolates will affect clinical efficacy, but they have emerged in a real world situation suggesting there is a significant benefit which has been selected in practice. Interestingly, octenidine dihydrochloride was only in use for one year before there was a marked change in the susceptibility of isolates.

For both biocides, isolates with decreased susceptibility were not clonal which suggests there has been no emergence of a dominant clone which may have an advantage in the face of either biocide. The fact that isolates with decreased susceptibility are seen in various strain backgrounds suggests that the capacity to develop this phenotype is common to many strains and, presumably results from a change in the *S. aureus* core genome. This differs to previous reports where there has been spread of a dominant clone with reduced susceptibility (Otter *et al*., 2013).

As highlighted by the review by Harbarth and colleagues the increased usage of antiseptics has not been matched by an increase in surveillance, both microbiological and clinical (Harbarth et al., 2014). One of the reasons for the lack of surveillance is the lack of both standardised testing methodology and definitions for resistance. The majority of studies have utilised MICs, but the methodologies vary and the technique is less applicable for antiseptics where the lethal rather than the inhibitory concentration needs to be measured and MBC testing is more meaningful. As there is no standardised testing methodology there has been no national or international agreement regarding an appropriate cut off for defining resistance. The clinical impact of reduced susceptibility demonstrated *in vitro* also needs to be assessed, with the concentrations of antiseptics measured *in vitro* are much lower than concentrations of antiseptics achieved *in vivo*.

Epidemiological typing in this study revealed that isolates with high MICs are spread across VNTR profiles, both in the predominant epidemic clone observed in the UK, ST22 and in CC8. When this is compared with the presence of *qacA/B* there was no correlation with chlorhexidine susceptibility; the majority of *qacA/B* isolates were from CC8. This clonal complex contains ST239 which has previously been described as carrying the *qacA/B* genes and has been associated with clinical failure of chlorhexidine (Batra *et al*., 2010). Carriage of *qacA/B* was only seen in one ST22 isolate in our panel, which is the most predominant ST both within the study hospital and in the UK. Despite the lack of carriage of *qacA/B* in ST22, the number of isolates with raised MICs was the same as amongst those from the CC8 group, highlighting further the lack of correlation between the presence of *qacA/B* and reduced susceptibility to chlorhexidine.

Consistent with the lack of correlation of *qacAB* carriage and antiseptic susceptibility, a genomic analysis of a subset of strains revealed no common mobile element which associated with chlorhexidine susceptibility. In two pairs of isolates with different chlorhexidine MICs mutations within chromosomal multidrug efflux systems *norA* or *norB* were found. NorA and NorB have previously been associated with chlorhexidine tolerance with isolates that over-express these systems demonstrating decreased chlorhexidine susceptibility (Liu et al., 2015; Vali et al., 2008, DeMarco et al., 2007) although another study reported only a weak association (Furi et al., 2013). Both *norA* and *norB* are part of the core *S. aureus* genome which is consistent with our observation that decreased susceptibility to chlorhexidine can emerge from multiple lineages and is not conditional on horizontal acquisition of *qacA/B*. Previous studies have focused on expression changes of efflux systems as a mechanism to increase efficiency of export of substrates. Mutation within the structural pump protein of a multidrug efflux system was also recently shown to increase efficiency of export of some substrates at the expense of others (Blair *et al*., 2015). The substitutions within NorB (adjacent to the translocation pore) and NorA (known to alter pump efficiency for fluoroquinolone export) identified here may reflect adaptation to increase efficiency of chlorhexidine export.

There is an obvious need for more research in this area to provide better surveillance data from larger populations and geographical regimes, and to understand the mechanisms of action and resistance to antiseptics better. The significance and clinical impact of the emergence of isolates with decreased susceptibility to decolonisation regimes remains uncertain. However, the introduction of chlorhexidine and octenidine dihydrochloride has changed the susceptibility of the *S. aureus* population compared to pre-biocide times and they are therefore having an ecological impact. The consequences of this remain unclear.

